# Improved SNV discovery in barcode-stratified scRNA-seq alignments

**DOI:** 10.1101/2021.06.12.448184

**Authors:** NM Prashant, Hongyu Liu, Christian Dillard, Helen Ibeawuchi, Turkey Alsaeedy, Kwan Hang Chan, Anelia Horvath

## Abstract

Single cell SNV analysis is an emerging and promising strategy to connect cell-level genetic variation to cell phenotypes. At the present, SNV detection from 10x Genomics scRNA-seq data is typically performed on the pooled sequencing reads across all cells in a sample. Here, we assess the gain of information of SNV assessments from individual cell scRNA-seq data, where the alignments are split by barcode prior to the variant call. For our analyses we use publicly available sequencing data on the human breast cancer cell line MCF7 cell line generated at consequent time-points during anti-cancer treatment. We analyzed SNV calls by three popular variant callers – GATK, Strelka2 and Mu-tect2, in combination with a method for cell-level tabulation of the sequencing read counts bearing SNV alleles – SCReadCounts. Our analysis shows that variant calls on individual cell alignments identify at least two-fold higher number of SNVs as compared to the pooled scRNA-seq. We demonstrate that scSNVs exclusively called in the single cell alignments (scSNVs) are substantially enriched in novel genetic variants and in coding functional annotations, in particular, stop-codon and missense substitutions. Furthermore, we find that the expression of some scSNVs correlates with the expression of their harbouring gene (cis-scReQTLs).

Overall, our study indicates an immense potential of SNV calls from individual cell scRNA-seq data and emphasizes on the need of cell-level variant detection approaches and tools. Given the growing accumulation of scRNA-seq datasets, cell-level variant assessments are likely to significantly contribute to the understanding of the cellular heterogeneity and the relationship between genetics variants and functional phenotypes. In addition, cell-level variant assessments from scRNA-seq can be highly informative in cancer where they can help elucidate somatic mutations evolution and functionality.

## Introduction

In single-cell studies, SNV analysis is an emerging and promising strategy to connect cell-level genetic variation to phenotypes and to interrogate lineage relationships in heterogeneous cell populations. To detect single-cell SNVs from DNA, genome and exome sequencing experiments can be performed [1–5]. These studies have revealed enormous amounts of knowledge on the cell-level genetic heterogeneity; however, they face challenges related to sample availability, unequal coverage and amplification bias, and are relatively costly for large scale applications. Recently, SNV detection from single cell RNA sequencing (scRNA-seq) experiments have started to emerge [6–9]. These analyses can complement DNA-based SNV-studies and maximize the potential of scRNA-seq datasets. Importantly, SNVs from scRNA-seq studies can provide crucial information on the SNV functionality through studying the allele specific dynamics and its correlation to phenotype features, such as gene expression and splicing[10– 12].

Among the scRNA-seq platforms, droplet-based technologies, such as 10x Genomics Chromium Single Cell 3′ and 5′ Expression workflows are quickly gaining popularity. Presently, SNV detection from 10x scRNA-seq data is typically performed on the pooled sequencing reads (pseudo-bulk), where it utilizes approaches optimized for bulk DNA-and RNA-variant calling [7,9,10,13] These approaches often estimate quality control metrics, such as variant allele fraction (VAF) and/or genotype confidence, based on all sequencing reads in a sample [14,15]. As a result, SNVs with low VAF and/or uncertain genotypes in the pooled data are frequently filtered out. While it is well acknowledged that post-zygotically occurring SNVs (such as somatic and mosaic mutations), being present in only a proportion of cells, can result in low VAF and uncertain genotypes, distinguishing those mutations from noise is difficult. Current approaches target somatic mutations by adjusting thresholds for VAF- and genotype-based filtering and accounting for population SNV frequencies [16]. Nevertheless, without cell-level information, detection of low-frequency SNVs is challenging. More recently, methods utilizing cellular barcodes for improved SNV detection have started to emerge [17,18].

In this study, we systematically assess the gain of information of SNV detection at each cell individually, where the alignments are split by barcode prior to the variant call. We reasoned that such setting enables VAF and genotype assessments per cell (as opposed to per sample), and consequently, is likely to result in retaining of additional high quality SNVs by variant callers.

To compare SNV assessments from single cells to those from pooled and bulk datasets, we utilized matched genome, exome, and scRNA-seq data from multiple time-points during anticancer treatment. We assessed SNV calls by three popular variant callers – GATK, Strelka2 and Mutect2 [14–16], in combination with a recently developed in our lab method for cell level tabulation of the sequencing read counts bearing SNV alleles from barcoded scRNA-seq alignments – SCReadCounts [18]. We then analyzed the SNVs called exclusively in the single cell alignments (single cell exclusive SNVs, scSNVs) for cellular distribution and annotations and correlated their allele expression to the expression level of their harboring gene.

Our results show that even after high stringency filtering, in the individual cell scRNA-seq data we identify at least two-fold higher number of SNVs, as compared to the unfiltered union of SNVs called in the pooled scRNA-seq, exome and genome sequencing data. We demonstrate that scSNVs exclusively called in the individual alignments (single cell SNVs, scSNVs) are substantially enriched in novel genetic variants and in coding functional annotations. Furthermore, we visualize the dynamics of the expression of these SNVs across multiple time-points during anti-cancer treatment and demonstrate correlations of the expression of some of these SNVs to the expression of their harboring gene. Overall, our analysis indicates an immense potential of SNV detection from individual cell scRNA-seq data and emphasizes on the need for cell-level variant assessment tools.

## Results

### Analytical pipeline

For our analyses we utilized publicly available scRNA-seq datasets of the breast cancer cell line MCF7 collected at four different time points during treatment with the anticancer agent bortezomib (Selleckchem, S1013) [19]. The four time-point sampling was performed before treatment (t0), after 12h (t12), 48h (t48), and 72h of exposure followed by drug wash and 24h of recovery (t96). These datasets were accompanied by matched whole genome sequencing (WGS) and deep (approximately 250× coverage) targeted exon sequencing (TES) of 334 genes that are commonly mutated in cancer (Profile OncoPanel v.3). We reasoned that the described data collection maximizes the identifiable SNVs across compatible DNA/RNA regions in bulk/pooled data. We then asked if additional SNVs can be identified using the barcode-stratified individual cell scRNA-seq data.

Our approach is presented on Figure 1. We first aligned the raw sequencing reads to GRCh38 using BWA for the DNA and STARsolo for the RNA data (Methods). For the initial SNV assessments we selected two widely used variant callers – GATK and Strelka2 - which have repeatedly demonstrated high quality performance across both DNA and RNA sequencing data, including scRNA-seq data [6,7,20]. For the bulk/pooled data we aimed to retain maximum number of identifiable SNVs. Accordingly, we applied GATK and Strelka2 in parallel, and then generated the union of the SNV calls across the genome, targeted exome, and each of the 4 corresponding RNA-sequencing datasets. To retain SNV calls with low VAF, the variant calls were not filtered for depth of allele coverage or confidence of the genotype call.

**Figure 1.**
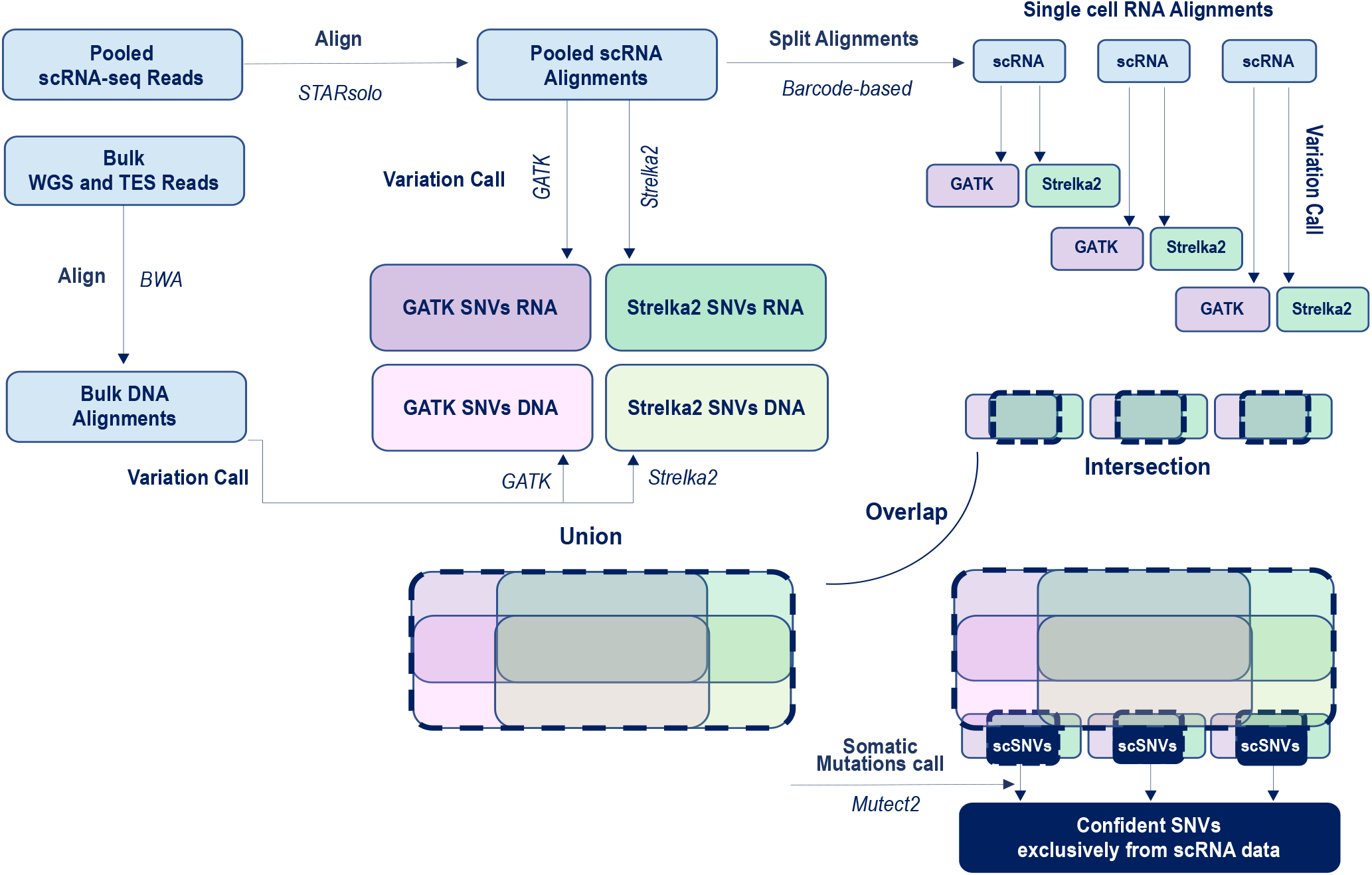
Analytical workflow for identification of confident SNVs calls exclusively in the individual scRNA-seq alignments, and not in any of the bulk or pooled sequencing datasets from the same sample.

Next, using the cellular barcodes and an in-house optimized script [21], we split the pooled scRNA-seq alignments into individual cell alignments each containing reads from a single cell. On each of these alignments we then applied GATK and Strelka2 in parallel, filtered the variant lists to retain only high-quality calls (Methods), and generated the intersection between GATK and Strelka2 to be used for our downstream analyses. Note that in contrast to the pool/bulk data, where we aim at maximizing the SNVs detection by combining SNVs detected by any of the callers, in the individual alignments our aim is to outline the highest confidence SNVs, for which we retain only high quality calls by both GATK and Strelka2.

### SNV calls across TES, WGS, and scRNA-seq

For this analysis, we applied the above described pipeline on the genomic regions compatible across TES, WGS and RNA-seq, which comprised the exons of the genes targeted by the POP exome capture. The numbers of common and exclusive SNVs in TES, WGS and pooled and individual scRNA-seq alignments are shown on Table 1 and S_Figure 1. In the individual alignments, Strelka2 called 2-to 3-fold higher number of SNVs, which included the vast majority of the GATK calls (Figure 2a).

**Table 1.**
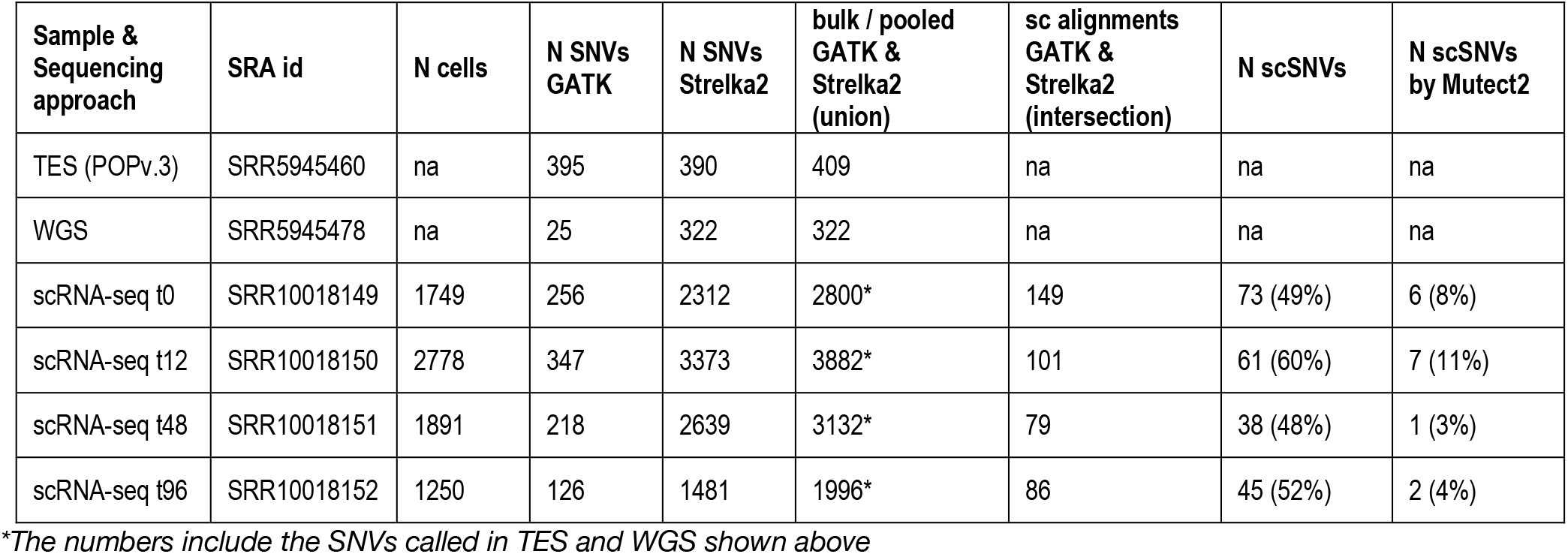
Number of SNVs identified in the MCF7 sequencing datasets in the exonic regions of the genes included in the ProfileOncoPanel (POPv.3)

| Sample & Sequencing approach | SRA id | N cells | N SNVs GATK | N SNVs Strelka2 | bulk / pooled GATK & Strelka2 (union) | sc alignments GATK & Strelka2 (intersection) | N scSNVs | N scSNVs by Mutect2 |
| --- | --- | --- | --- | --- | --- | --- | --- | --- |
| TES (POPv.3) | SRR5945460 | na | 395 | 390 | 409 | na | na | na |
| WGS | SRR5945478 | na | 25 | 322 | 322 | na | na | na |
| scRNA-seq t0 | SRR10018149 | 1749 | 256 | 2312 | 2800* | 149 | 73 (49%) | 6 (8%) |
| scRNA-seq t12 | SRR10018150 | 2778 | 347 | 3373 | 3882* | 101 | 61 (60%) | 7 (11%) |
| scRNA-seq t48 | SRR10018151 | 1891 | 218 | 2639 | 3132* | 79 | 38 (48%) | 1 (3%) |
| scRNA-seq t96 | SRR10018152 | 1250 | 126 | 1481 | 1996* | 86 | 45 (52%) | 2 (4%) |

\*The numbers include the SNVs called in TES and WGS shown above

**Figure 2.**
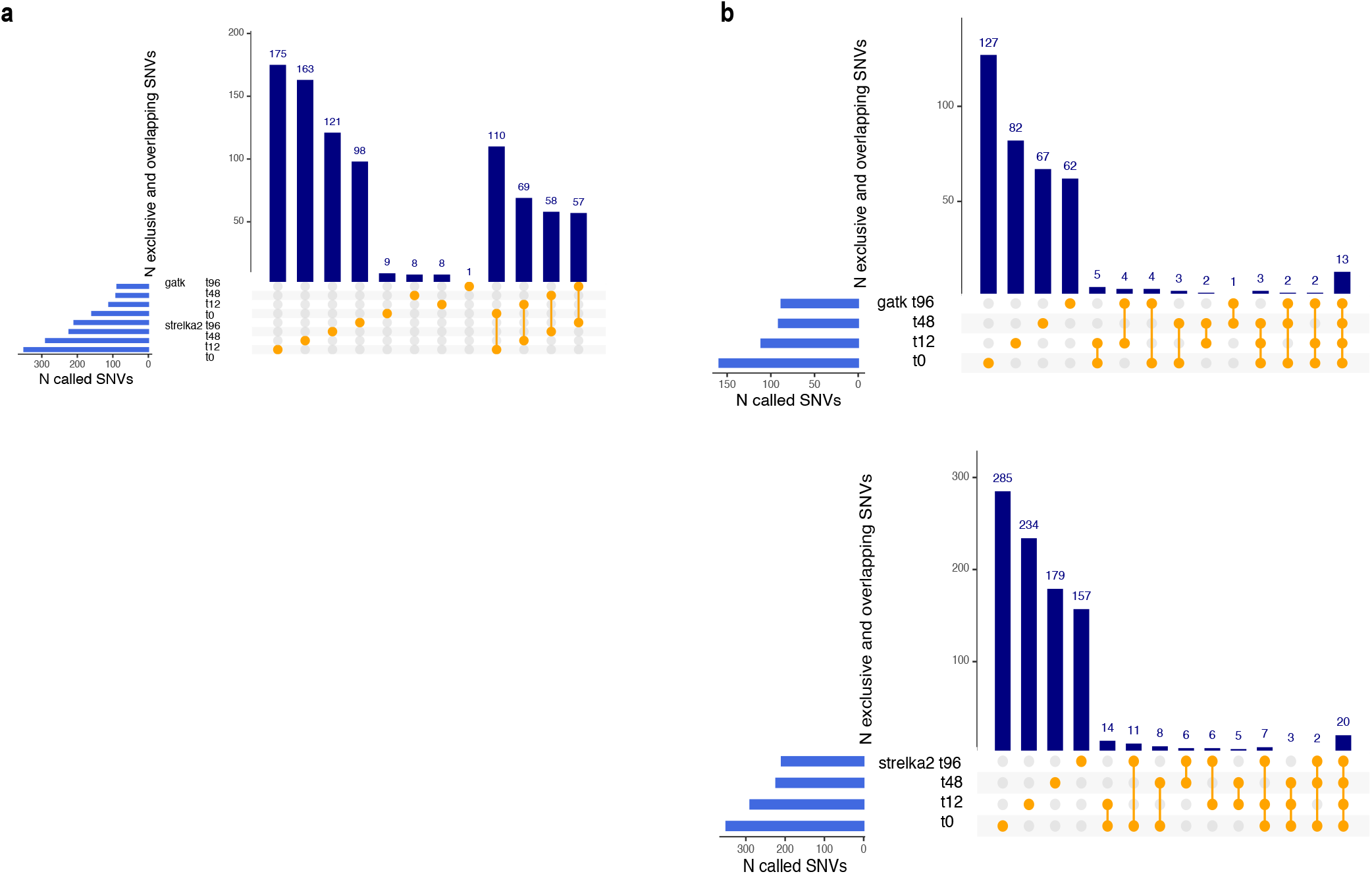
**a.** Concordance between GATK and Strelka2 in variant calling from individual cell alignments. Higher number of SNVs are called by Strelka2, which also identifies the vast majority of the GATK calls. Note that the UpSet plots show the first 12 of all possible overlaps. **b**. Shared and exclusive scSNVs called by GATK (top) and Strelka2 (bottom) from scRNA-seq data generated at four time-points during drug treatment show low overlap indicative for de novo SNVs.

Across the four scRNA-seq datasets, in the exonic regions of the 334 genes from the POP panel, the above pipeline identified between 38 and 73 scSNVs (S_Table 1; all of these SNVs were visually examined through the Integrative Genome Viewer. These numbers represented 48% and above of all confident individual alignment SNVs. To assess if the scSNVs are identifiable with callers specifically targeting SNVs in low proportion of cells, we applied Mutect2 [16]. In the pooled data, Mutect2 identified up to 11% of the scSNVs. Thus, our analysis shows that even in settings strongly favouring variant discovery from bulk/pooled data, assessments of barcode-stratified individual cell alignments detect substantially higher number of SNVs.

We next assessed the proportion of SNVs shared across the four time-points post drug treatment. This analysis was performed separately for the GATK and Strelka2, which showed highly concordant results. As seen on Figure 2b, scSNVs show low overlap across the samples collected over the four time-points of drug treatment. This suggests enrichment with *de novo* arising SNVs, which is consistent with the finding of the original study on quick MCF7 evolution during anticancer treatment [19].

### Transcriptome-wide SNVs called exclusively in the individual alignments

Following the above described strategy we next analysed the transcriptome-wise shared and exclusive SNVs between the pooled and scRNA-seq alignments of the 4 time-points; the results are summarized in Table 2. Specifically, in the individual alignments, after stringent filtering of both GATK and Strelka2 calls, and retaining only the intersection of the two callers, we identified between 7 and 14 thousands SNVs per dataset that were not captured in the pooled scRNA-seq data by neither GATK or Strelka2 (S_Table 2). From those, only up to 10% were identified using Mutect2. This observation is aligned with the findings across the WGS/TES/RNA datasets on the exons of the POP capture and suggests that transcriptome-wise application of variation call on barcode-stratified individual scRNA-seq data can identify thousands of SNVs in addition to those identified in the pooled scRNA-seq data.

**Table 2.** Number of SNVs transcriptome-wise, and percent cells with scSNVs.

| Sample | Pooled scRNA-se alignments<br>(GATK & Strelka2 union) | N scSNVs<br>(GATK&Strelka2<br>intersection) | N scSNVs<br>by Mutect2 | Max Percent<br>cells with<br>scSNVs | N scSNVs<br>in 2 and more<br>cells |
| --- | --- | --- | --- | --- | --- |
| scRNA-seq t0 | 489048 | 13385 | 1310 (9.8%) | 90 / 1749 (5,2%) | 636 (4,8%) |
| scRNA-seq t12 | 524598 | 9470 | 936 (9.9%) | 44 / 2778 (1,6%) | 472 (5%) |
| scRNA-seq t48 | 446779 | 7131 | 560 (7.8%) | 33 / 1891 (1,8%) | 318 (4.5%) |
| scRNA-seq t96 | 335839 | 10794 | 856 (7.9%) | 30 / 1250 (2,4%) | 429 (4%) |

We next assessed the number of cells bearing each of the scSNVs. As expected, the maximum percentage of cells with scSNVs represented up to 5% of the cells in the dataset (See Table 2), with the majority of the scSNVs seen in only one cell (S_Figure 2). Between 318 and 636 scSNVs (between 4 and 5% of all scSNVs per sample) were called in two or more cell (See S_Figure 2).

### Novel and known SNVs in the individual scRNA-seq alignments

We next analysed the proportion of novel (previously undescribed) scSNVs, and compared to the proportion on novel SNVs identified in the pooled scRNA-seq datasets (pSNVs). For this analysis, we used pSNV calls processed exactly the same way as the scSNVs (i.e. the intersection of filtered GATK and Strelka2 calls). As novel we assigned SNVs not present in the NCBI Single Nucleotide Polymorphism database (DbSNP), the Catalogue of Somatic Mutations in Cancer (COSMIC) or the ATLAS of RNA editing events in human (REDIportal) [22,23].

Notably, among the scSNVs we estimated several-folds higher proportion of novel SNVs. Specifically, over 70% of the scSNVs in each of the datasets were novel, compared to up to 15% novel SNVs called in the corresponding pooled scRNA-seq datasets (Figure 3a). This difference is likely due to the suggested high rate of *de novo* acquired mutations, present in small proportion of cells and therefore detectable exclusively in the individual scRNA-seq alignments.

**Figure 3.**
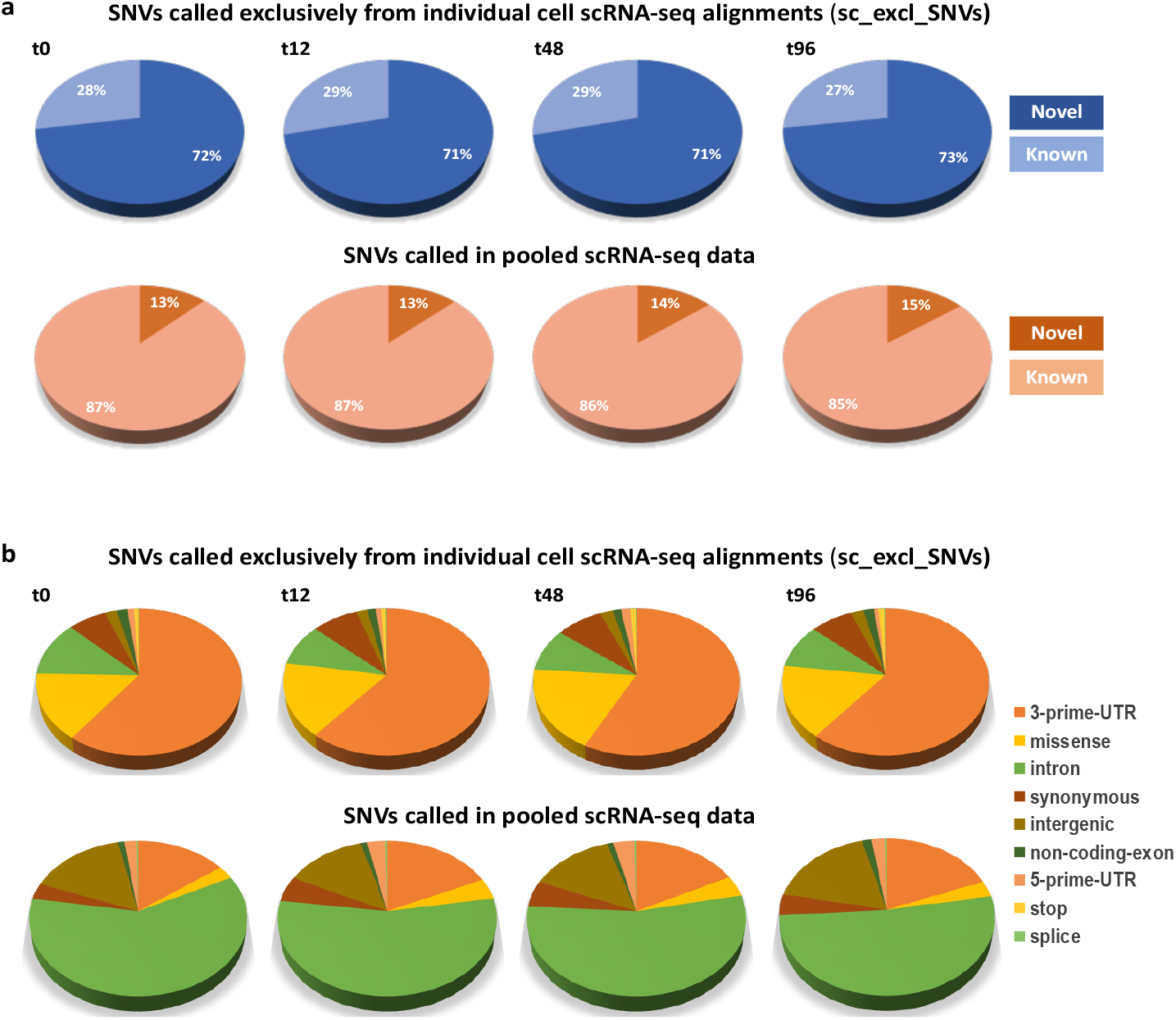
**a.** Percentage of novel and known SNVs called exclusively in the individual alignments (top) and in the pooled scRNAseq data (bottom). Approximately 5-fold higher percentage of novel SNVs is seen in the individual cell alignments. **b**. Distribution of functional annotations among the SNVs called exclusively in the individual alignments (top), as compared to the pooled scRNA-seq data (bottom). Significantly higher proportions of 3’-prime-UTR, missense and stop-codon SNVs are called in the individual alignments.

Next, we compared the distribution of predicted SNV functional annotations. This analysis revealed significant differences in the proportions of all the functional annotations between the scSNVs and the SNVs called in the pooled scRNA-seq (Figure 3b). The largest annotation category for the scSNVs was 3’-UTR, whereas for the SNVs in the pooled data it was intronic. ScSNVs also had a significantly higher proportion of coding variants, including stop-codon and missense substitutions (Table 3). The most striking difference was estimated for the stop codon mutations which showed approximately 50-fold higher rate among the scSNVs (around 1%, as opposed to up to 0.02% in the pooled SNVs). The missense substitutions had 4-to 6-fold higher rate among the scSNVs. In contrast, synonymous SNVs and SNVs in non-coding exons showed only up to 2-fold higher rate in the scSNVs.

**Table 3.** Distribution of functional annotations between scSNVs and pSNVs, chi-square comparisons

| Function | t0 |  | t12 |  | t48 |  | t96 |  |
| --- | --- | --- | --- | --- | --- | --- | --- | --- |
|  | sc_excl<br>SNVs | pool<br>SNVs | sc_excl<br>SNVs | pool<br>SNVs | sc_excl<br>SNVs | pool<br>SNVs | sc_excl<br>SNVs | pool<br>SNVs |
| 3-prime-UTR | 8075<br>(60%) | 6062<br>(15%) | 5785<br>(61%) | 6340<br>(18%) | 4109<br>(57%) | 5618<br>(17%) | 6562<br>(61%) | 5272<br>(19%) |
| chi_sq, p<br>P-value | 10883<br>p<10e-7 |  | 6981<br>p<10e-7 |  | 5092<br>p<10e-7 |  | 6403<br>p<10e-7 |  |
| missense | 2010<br>(15%) | 1129<br>(3%) | 1562<br>(17%) | 1513<br>(4%) | 1323<br>(18%) | 1388<br>(4%) | 1744<br>(16%) | 915<br>(3%) |
| chi-square<br>P-value | 2808<br>p<10e-7 |  | 1726<br>p<10e-7 |  | 1854<br>p<10e-7 |  | 2008<br>p<10e-7 |  |
| intron | 1609<br>(12%) | 24634<br>(60%) | 870<br>(9%) | 19247<br>(55%) | 667<br>(9%) | 17559<br>(54%) | 1045<br>(10%) | 14351<br>(52%) |
| chi-square<br>P-value | 9236<br>p<10e-7 |  | 6261<br>p<10e-7 |  | 4749<br>p<10e-7 |  | 5694<br>p<10e-7 |  |
| synonymous | 897<br>(7%) | 1365<br>(3%) | 758<br>(8%) | 1844<br>(5%) | 580<br>(8%) | 1756<br>(5%) | 793<br>(7%) | 1213<br>(4%) |
| chi-square<br>P-value | 287<br>p<10e-7 |  | 102<br>p<10e-7 |  | 76<br>p<10e-7 |  | 140<br>p<10e-7 |  |
| intergenic | 276<br>(2%) | 6484<br>(16%) | 182<br>(2%) | 4680<br>(13%) | 161<br>(2%) | 4475<br>(14%) | 236<br>(2%) | 4941<br>(18%) |
| chi-square<br>P-value | 1751<br>p<10e-7 |  | 997<br>p<10e-7 |  | 755<br>p<10e-7 |  | 1624<br>p<10e-7 |  |
| non-coding-<br>exon | 250<br>(1,9%) | 414<br>(1%) | 142<br>(1,5%) | 357<br>(1%) | 102<br>(1,4%) | 331<br>(1%) | 200<br>(1,9%) | 421<br>(1,5%) |
| chi-square<br>P-value | 60<br>p<10e-7 |  | 15<br>p=0.00007 |  | 9<br>p=0.002 |  | 6<br>p=0.02 |  |
| 5-prime-UTR | 152<br>(1,1%) | 816<br>(2%) | 78<br>(0,8%) | 1027<br>(2,9%) | 118<br>(1,6%) | 1098<br>(3,4%) | 80<br>(0,7%) | 622<br>(2,2%) |
| chi-square<br>P-value | 41<br>p<10e-7 |  | 135<br>p<10e-7 |  | 58<br>p<10e-7 |  | 96<br>p<10e-7 |  |
| splice | 16<br>(0,12%) | 105<br>(0,26%) | 15<br>(0,16%) | 120<br>(0,34%) | 9<br>(0,13%) | 103<br>(0,32%) | 18<br>(0,17%) | 78<br>(0,28%) |
| chi-square<br>P-value | 8<br>p=0.005 |  | 8<br>p=0.006 |  | 7<br>p=0.008 |  | 4<br>p=0.04 |  |
| stop | 100<br>(0,75%) | 6<br>(0,01%) | 78<br>(0,82%) | 6<br>(0,02%) | 62<br>(0,86%) | 2<br>(0,01%) | 116<br>(1,07%) | 5<br>(0,02%) |
| chi-square<br>P-value | 275<br>p<10e-7 |  | 253<br>p<10e-7 |  | 264<br>p<10e-7 |  | 275<br>p<10e-7 |  |

The observed differences in the functional categories in the scSNVs requires further attention and analyses on high number of samples. Like the high proportion of novel mutations, it is likely to be related to *de novo* scSNVs, where different rates of mutations generation, mismatch repair and purifying selection across different functional genomic regions play role. Nevertheless, our observation highlights the potential of the scRNA-seq analyses to study mutation dynamics and evolution.

### ScSNVs expression

To estimate the expression of the scSNVs we applied SCReadCounts as previously described [18]. For each cell, SCReadCounts tabulates the reference and variant counts of sequencing reads (nref and n_var_, respectively) for genomic positions of interest, and computes the expressed Variant Allele Fraction (VAF_RNA_ = n_var_ / (n_var_ + n_ref_)) at a desired depth threshold (minimum number of reads covering the position, minR). For this particular analysis we estimated VAF_RNA_ at minR=3. The distribution of VAF_RNA_ for scSNVs called in 3 and more cells per dataset, for all cells with 3 and more reads at the corresponding position, is shown on Figure 4a. The majority of the scSNV positions had VAF_RNA_ up to 0.2 across most of the cells. Note that this assessment includes also the cells with only reference reads at the SNV position (i.e. VAF_RNA_ = 0). Such VAF_RNA_ distribution is expected for SNVs present in a small proportion of cells (i.e. *de novo* SNVs). In contrast, biallelic pSNVs show VAF_RNA_ distribution centered around 0.5, which is generally expected for the majority of the heterozygous germline SNVs (Figure 4b).

**Figure 4.**
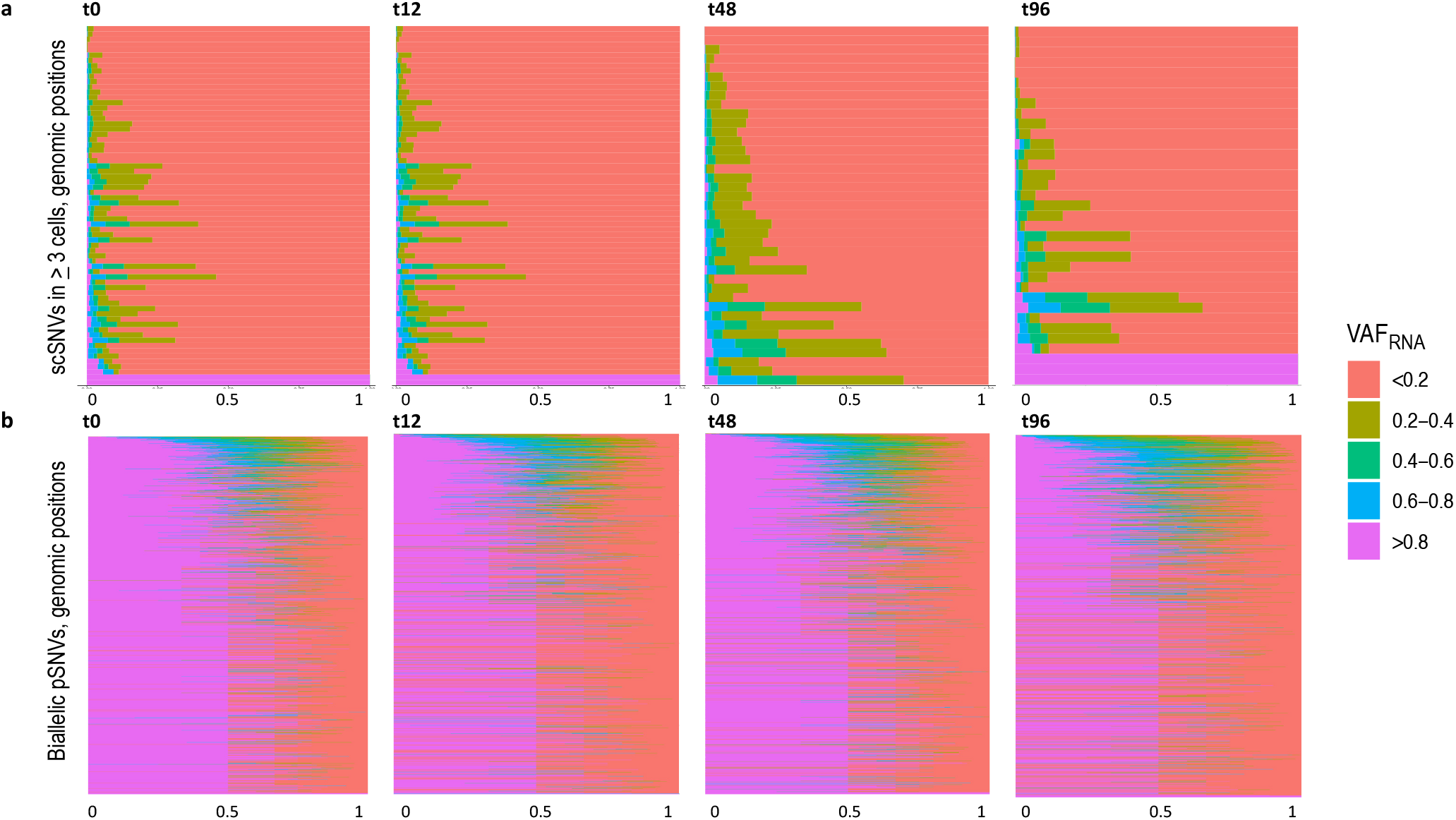
**a.** ScVAF_RNA_ estimated at positions covered by a minimum of 3 sequencing reads (minR=3) for scSNVs called in 3 and more cells per dataset (y-axis). The majority of the positions have VAF_RNA_ up to 0.2. Note that the plot is inclusive for all the cells with minR=3 in the corresponding position, including those covered with reference reads only. The percentage of cells with the corresponding VAF_RNA_ is displayed on the x-axis **b**. ScVAF_RNA_ estimated at positions covered by a minimum of 3 sequencing reads for biallelic pSNVs (y-axis). For most of the pSNVs VAF_RNA_ distribution is centered around 0.5, which is expected for germline heterozygous SNVs not subjected to monoallelic expression.

To explore if cells bearing certain scSNVs have related gene expression features, we assessed the scSNV expression in the individual cells after graph-based cell-clustering. For this analysis we processed the scRNA-seq datasets as we have previously described [11,18]. Briefly, after alignment with STARsolo [12] and quality filtering, the gene-expression matrices were processed using Seurat [24] to normalize gene expression and correct for batch- and cell-cycle effects; the normalized gene expression values were then used to assign likely cell types using SingleR [25] (Methods). We then visualized VAF_RNA_ in the cells bearing scSNVs over the UMAP two-dimensional projections of the scRNA-seq datasets; examples are shown on Figure 5.

**Figure 5.**
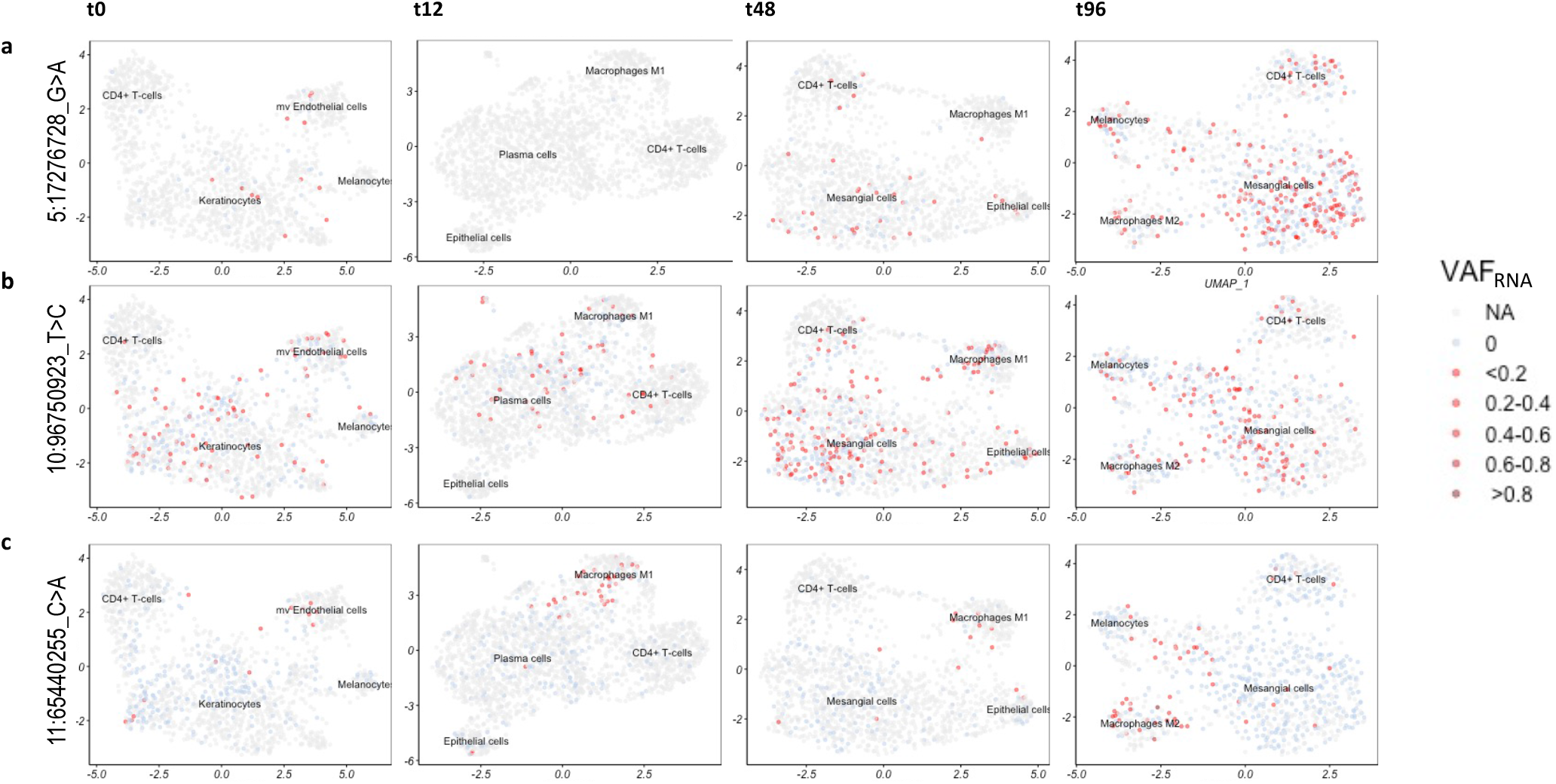
Two-dimensional UMAP projections with quantitative visualization (red) of scSNVs VAF_RNA_. Light-blue colour indicates that the position is covered by at least 3 unique sequencing reads bearing the reference nucleotide; thereby signifying non-0 expression at the position. **a.** SNV rs1161976348 (5:17276721_G>A) in the 3’-UTR of the gene *BASP1*. Higher proportion of cells appear to express the SNV at later time-points post anti-cancer treatment, especially at t96. **b**. Novel intergenic SNV (10:96750923_T>C) showing relatively even distribution across the different cell types and clusters of the 4 post-treatment time-points. **c**. Novel SNV positioned at 11:65440255 (C>A) in a non-coding exon of the gene *NEAT1*, expressed preferentially in the Microphages.

Some scSNVs showed different expression across the four treatment time-points. An example is rs1161976348 (5:17276721_G>A in the 3’-UTR of the gene *BASP1*), which appeared to be expressed in a higher proportion of cells at later time-points, and especially at t96 (Figure 5a). Other scSNVs (such as the novel intergenic SNV 10:96750923_T>C) showed relatively even distribution across the different cell types and clusters (Figure 5b). In contrast, the novel SNV positioned at 11:65440255_C>A in a non-coding exon of the gene *NEAT1* showed preferential expression in Macrophage-like cells (Figure 5c).

Finally, we assessed if some of the scSNVs are correlated the expression of their harboring gene, for which we applied the linear regression model implemented in scReQTL [11]. For this analysis we used scSNVs detected in five and more cells (between 35 and 70 scSNVs per datasets). Across the four datasets, we identified a total of 20 cis-scReQTLs at significance level p<0.05 (Figure 6 and S_Figure 3).

**Figure 6.**
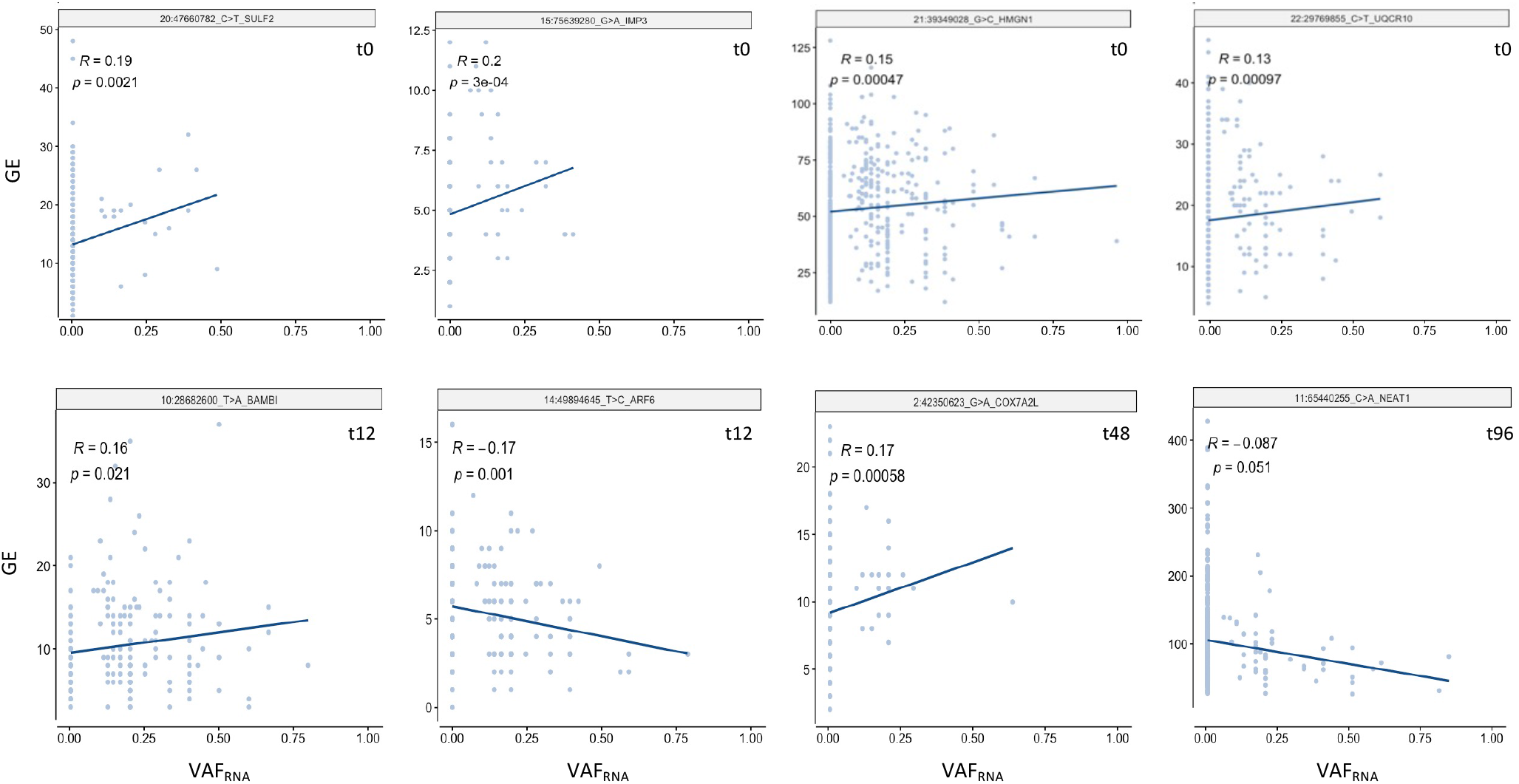
Significant cis-scReQTLs at FDR<0.05.

## Discussion

In this study we perform an initial assessment SNV calls for individual barcode stratified scRNA-seq alignments. Our analysis shows that this strategy identifies significantly higher number of SNVs as compared to variant calls on pooled/bulk scRNA-seq data. Furthermore, we find that SNVs called exclusively in the individual alignments (and not in the pooled/bulk scRNA-seq data) possess several striking characteristics. First, scSNVs are substantially enriched in previously undescribed variants. This finding is not surprising, as scSNVs are seen in up to 5% of the cells in a datasets (most often in only one cell) and thus are likely to indicate *de novo* arising variants. *De novo* SNVs are likely to arise in most of the normal and tumor cells [26], but are only possible to be retained in the germline in germline tissues. Therefore, scSNVs are unlikely to be reported in DbSNP, where the vast majority of SNVs are called from pooled germline DNA datasets. Hence, calling scSNVs can facilitate studies on the occurrence and the evolution of *de novo* genetic variants. We note that here we cannot exclude the possibility for RNA-editing origin of some of the SNVs. However, we find the probability for RNA-editing as low since none of these loci was listed in RNA-editing databases, and, also because we remove from our analysis repeated regions (Methods), which are known to harbour the vast majority of RNA-editing events.

Second, we find that the scSNVs are significantly enriched in coding variants, and especially in stop-codon and missense substitutions. This is likely to be related to different rates of mutations’ generation, repair, and positive or negative selection. This observation definitely requires scRNA-seq studies focused on mutation dynamics and evolution, and exploiting multiple heterogeneous sample sources.

Third, we find that some scSNVs might affect the expression of their harbouring gene, and thereby, possibly exert downstream effects. In this study we find 20 significant cis-scReQTLs. This number is expected given the input size (up to 70 SNVs and up to 3000 cells per dataset), and, based on our previous studies is likely to be significantly higher in larger datasets [11,18].

Overall, our study indicates an immense potential of SNV assessment from individual cell scRNA-seq data and emphasizes on the need of cell-level variant assessment approaches and tools. Given the growing accumulation of scRNA-seq datasets, cell-level variant assessment are likely to significantly contribute to the understanding of the cellular heterogeneity and the relationship between genetics and functional phenotypes. Finally, cell-level variant assessments from scRNA-seq can be highly informative in cancer where they can help elucidate somatic mutations evolution and functionality.

## Methods

### Sequencing datasets

The sequencing datasets were freely available from the NCBI Sequence Read Archive (SRA) under the accession numbers SRR5945460 (MCF7, targeted exome), SRR5945478 (MCF7, whole genome), and SRR10018149, SRR10018150, SRR10018151, SRR10018152 (MCF7 treatment, t0, t12, t48 and t96, respectively). MCF7 cells culturing and treatment is described in details in the original study [19]. Briefly, to follow transcriptional changes during treatment, MCF7 cells were exposed to 500nM of bortezomib (Selleckchem, S1013) and collected before treatment (t0), after 12h of exposure (t12), after 48h of exposure (t48) or after 72h of exposure followed by drug wash and 24h of recovery (t96). Single cells were processed through the Chromium Single Cell 3′ Solution platform using the Chromium Single Cell 3′ Gel Bead, Chip and Library Kits (10X Genomics). Libraries were sequenced on an Illumina NextSeq 500 platform.

### Data processing

#### Alignment, barcode and UMI processing, generation of individual scRNA-seq alignments

The targeted exome and the whole genome sequencing reads were aligned to the latest version of the human genome reference (GRCh38, Dec 2013) using BWA v.0.7.17 default settings [27]. The pooled sequencing reads form the scRNA-seq datasets were aligned using the STARsolo module of STAR v.2.7.7a in 2-pass mode with transcript annotations from the assembly GRCh38.79 [12,28]. STARsolo integrates read mapping, read-to-gene assignment, cell barcode demultiplexing and unique molecular identifier (UMI) collapsing [12]. To generate individual cell alignments we adopted a publicly available python script which splits the pooled scRNA-seq alignments based on cellular barcode [21].

#### Variant call

For all DNA and RNA datasets variant call was performed applying the HaplotypeCaller module of GATK v.4.2.0.0, in parallel with Strelka2 v.2.9.10; both tools were used under their default settings [14,15]. For RNA datasets the HaplotypeCaller was preceded by assignment of read groups using the GATK module AddOrReplaceReadGroups, followed by splitting reads that contain Ns in their cigar string with the GATK module SplitNCigarReads [14]. For the initial comparisons that included the DNA datasets, no filtering was applied on the SNV calls from the pooled or bulked variant calls. The SNV calls from the individual alignments were filtered according to the following criteria using bcftools v.1.10.2 [29]: QUAL (Phred-scaled probability)> 100, MQ (mapping quality)> 60, and QD (quality by depth)>2. The same filtering was applied on the SNV calls from pooled alignments for the analyses of distribution on novel SNVs and functional annotations. SNV loci were annotated using SeattleSeq v.16.00 (dbSNP build 154), and SNV loci positioned in repetitive regions were removed. Thus processed SNV calls were subjected on the above described analyses.

#### Gene expression estimation from scRNA-seq data

To estimate gene expression, we used the read count matrices with the row gene counts per cell generated by STARsolo. We normalized and scaled the expression data using the sctransform function as implemented in Seurat v.3.0 [24,30], which stabilizes the gene expression variance using regularized negative binomial regression, and outlines the most variable genes. The sctransform function integrates the previous Seurat functions NormalizeData, ScaleData, and FindVariableFeatures. The cell-feature distributions were than plotted to identify and filter out outliers and low-quality cells, which we defined after examination of the cell features distribution (S_Figure 4). Specifically, based on the cells’ and features’ distribution, we have filtered out: (1) cells with mitochondrial gene expression over between 7.5% and 15%, (2) cells with less than 1000 genes, and (3) cells with more than between 4500 and 5500 detected genes (to remove potential doublets). The Seurat processed gene expression values were also used to remove batch effects and cell cycle effects (S_Figure 5), as well as for cell types assessments and cis-scReQTLs (See below).

#### Cell type assessments

To define similarity of the MCF7 clusters with known cell types, we used SingleR v.1.0.5 [25]. SingleR assigns cellular identity by comparison to reference whole transcriptome expression data sets of pure cell types. SingleR correlates the expression profile of each single cell to whole-transcriptome expression data from established cell types (BluePrint + ENCODE datasets). To select the expression profile most similar to the tested cells, the analysis is rerun iteratively, using only the top cell types from the previous step until only one cell type is retained. Comparing our datasets against 259 bulk RNAseq profiles representing 24 main cell types and 43 subtypes, SingleR identified the following cell types: CD4+ T-cells, Epithelial cells, Macrophages, Endothelial cells, Erythrocytes, Keratinocytes, Plasma cells, and Mesanglial cells (S_Figure 6).

#### VAF_RNA_ estimation

Single cell level VAF_RNA_ is assessed from the pooled scRNA-seq alignments using scReadCounts v.1.1.4 as we have previously described [18]. Briefly, provided barcoded scRNA-seq alignments and genomic loci and alleles of interest, SCReadCounts tabulates, for each cell, the reference and variant read counts (n_ref_ and n_var_, respectively), and generates a cell-SNV matrix with the absolute n_ref_ and n_var_ counts, and a cell-SNV matrix with the VAF_RNA_ estimated at a user-defined threshold of minimum number of required sequencing reads (minR). For the herein presented analysis, we used minR≥3.

#### Correlation between VAF_RNA_ and Gene Expression

For each scSNV called in more than 5 cells, we performed analysis for a correlation between the VAF_RNA_ and the gene expression (cis-scReQTL) of the harbouring gene using scReQTL as previously described [11]. Briefly, the VAF_RNA_ were correlated to the normalized gene expression values of the most variable genes using a linear regression model as implemented in Matrix eQTL [31]. The top 15 principal components of the gene expression were used as covariates. Cis-correlations were annotated as previously described for the bulk ReQTLs [32]. Briefly, because scReQTLs are assessed from transcripts, we assign cis-correlation based on the co-location of the SNV locus within the transcribed gene, using the gene coordinates.

#### Statistical analyses

Throughout the analysis we used the default statistical tests (with built-in multiple testing corrections) implemented in the used software packages (Seurat, SingleR, Matrix eQTL), where *p*-value of 0.05 was considered significant, unless otherwise stated. For estimation of significant scReQTL, we applied FDR as implemented in the Matrix eQTL package. Specifically, once Matrix eQTL discovers a set of significant gene-SNP pairs, it estimates a corresponding q-value (FDR) for each of them using Benja-mini–Hochberg procedure under the assumption that the tests are independent or positively correlated [31,33].

## Funding

This work was supported by MGPC, The George Washington University; [MGPC_PG2020to AH].

## Conflict of Interest

None declared.

## Ethics approval and consent to participate

The study uses only previously published and freely available datasets.

## Availability of data and materials

All the data analyzed in this study are supplied with the supplemental material or available as indicated in the cited publications.

## Authors’ contributions

PNM, HL, CD, HI, TA and HC performed the data processing and contributed to the analyses and the visualization; AH devised and supervised the study and wrote the manuscript.

## Notes

### Competing Interest Statement

The authors have declared no competing interest.

