## Supplementary material for "Improved SNV discovery in barcode-stratified scRNA-seq alignments": S_Figure 1

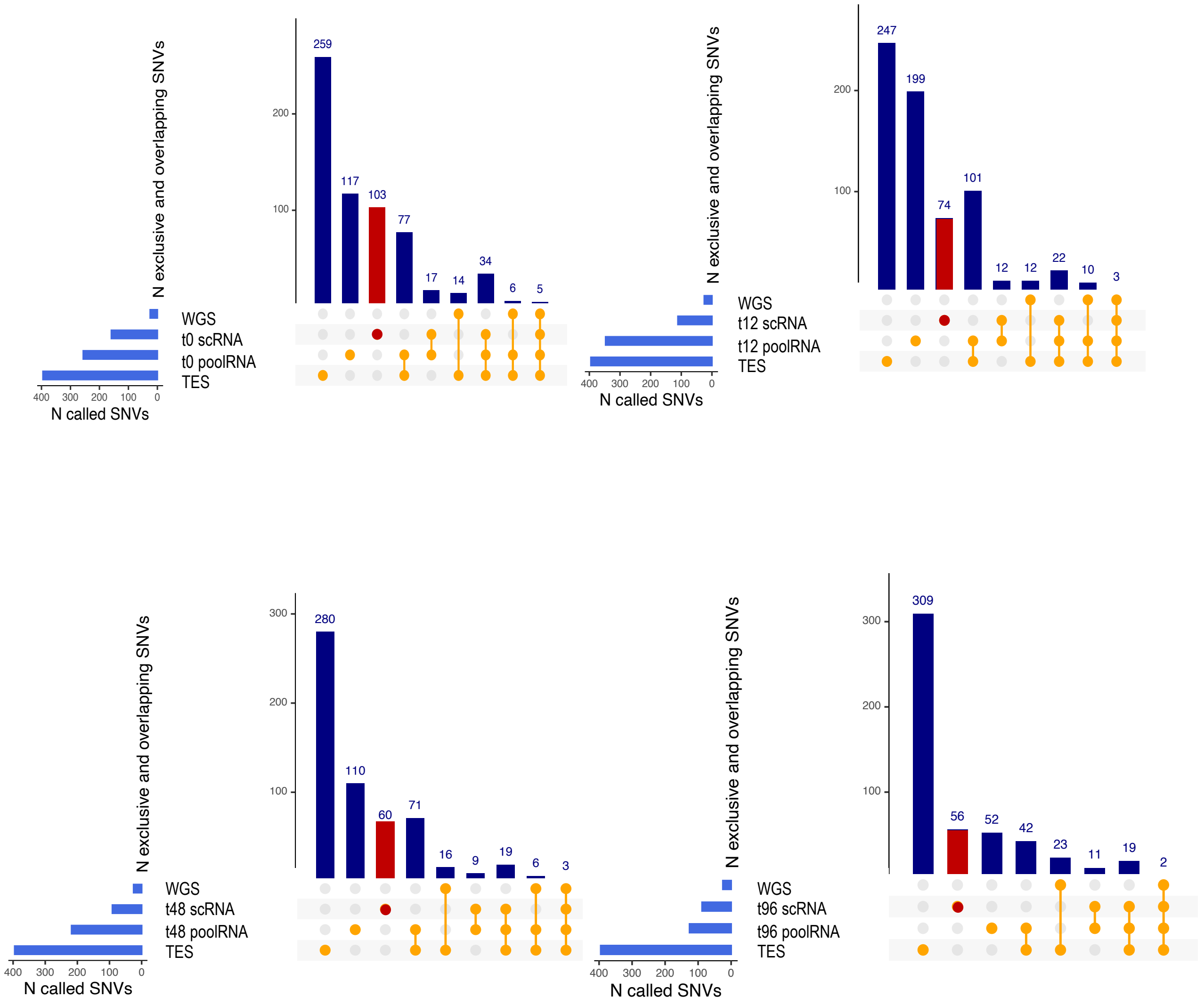


**S_Figure 1a.** Shared and exclusive SNVs calls across bulk WGS, TES, pooled scRNA-seq and individual cell alignments by GATK in the exons of genes included in the POP. The SNVs called exclusively in the single cell alignments are shown in red.

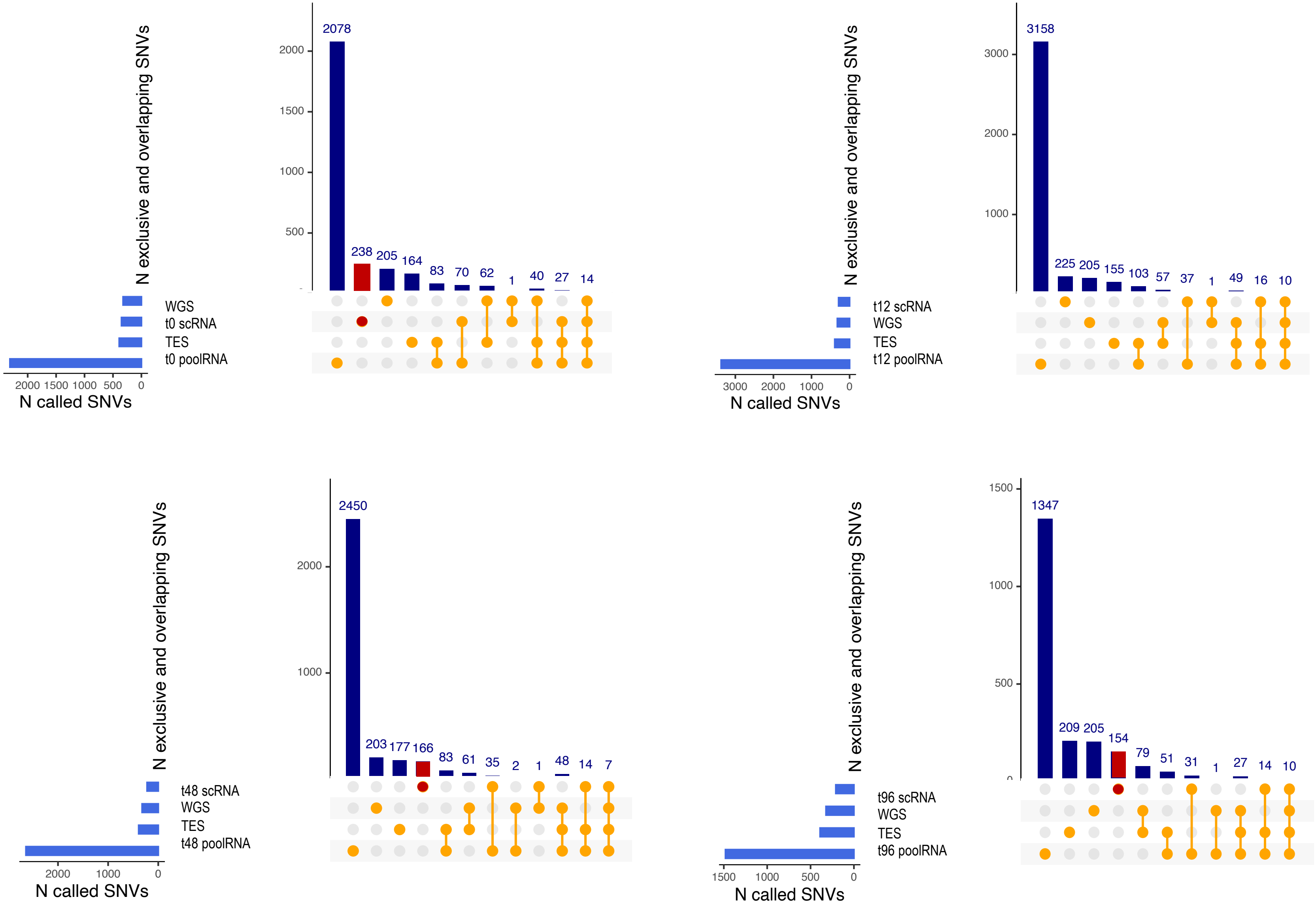


**S_Figure 1b.** Shared and exclusive SNVs calls across bulk WGS, TES, pooled scRNA-seq and individual cell alignments by Strelka2 in the exons of genes included in the POP. The SNVs called exclusively in the single cell alignments are shown in red.
