## Supplementary material for "Improved SNV discovery in barcode-stratified scRNA-seq alignments": S_Figure 2

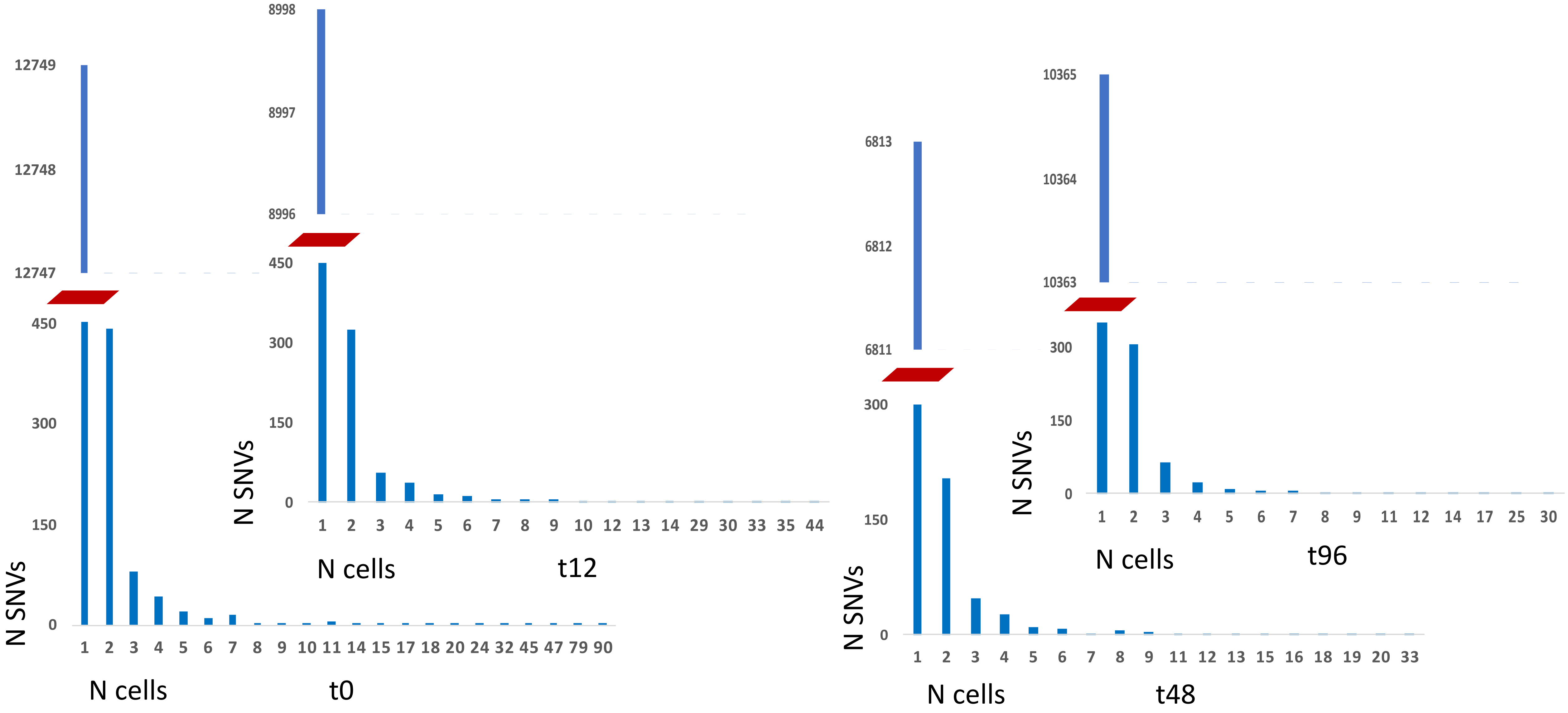


**S_Figure 2.** Cell distribution of sc_excl_SNVs across the 4 time-points of anticancer treatment of MCF7. Between 318 and 636 sc_excl_SNVs were called in more than one cell.
