## Supplementary figures and images for "Improved SNV discovery in barcode-stratified scRNA-seq alignments"

### S_Figure 3

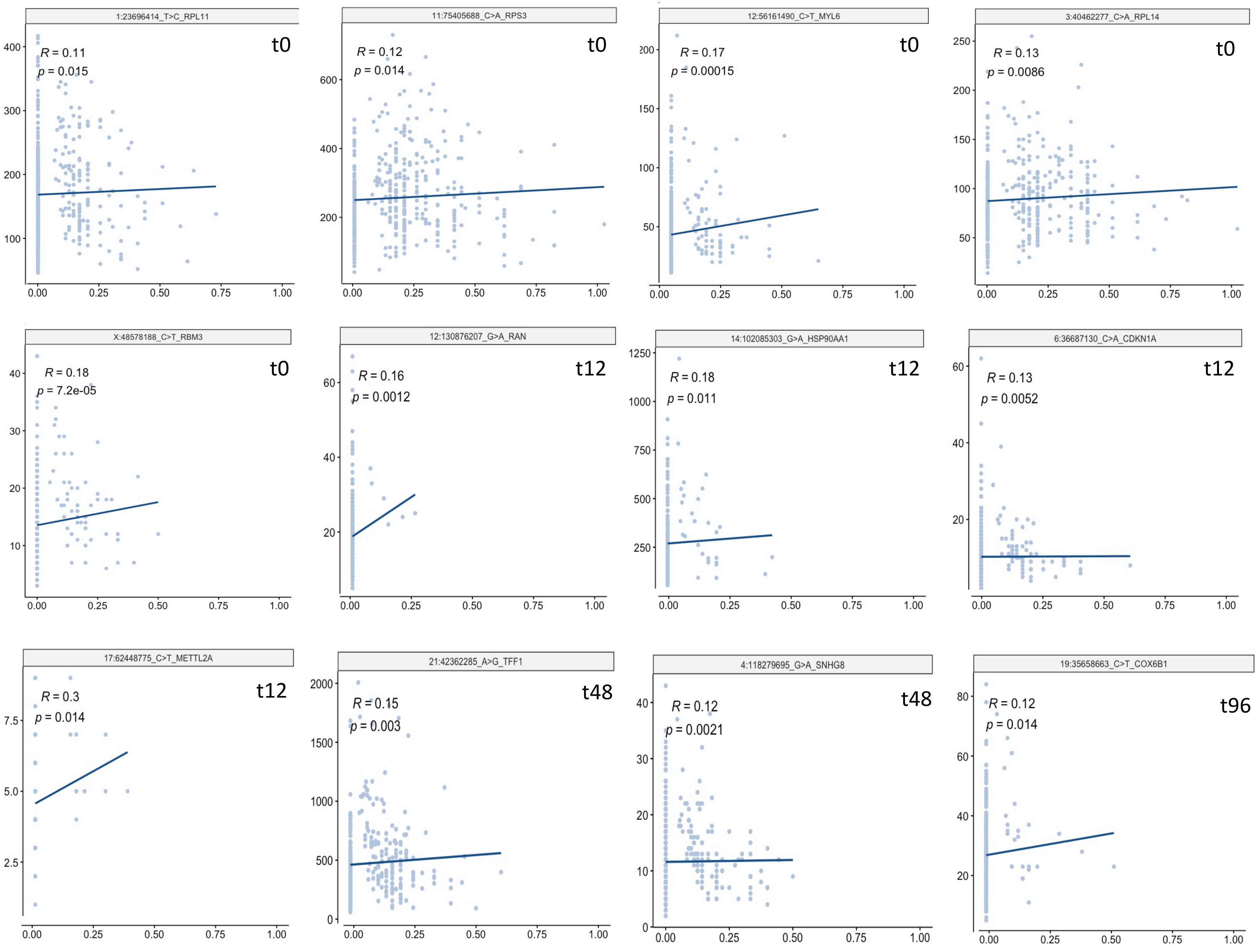


S_Figure 3 – signifcant cis-scReQTL between scSNVs and the expression of their harbouring gene.
