## Supplementary material for "Improved SNV discovery in barcode-stratified scRNA-seq alignments": S_Figure 4

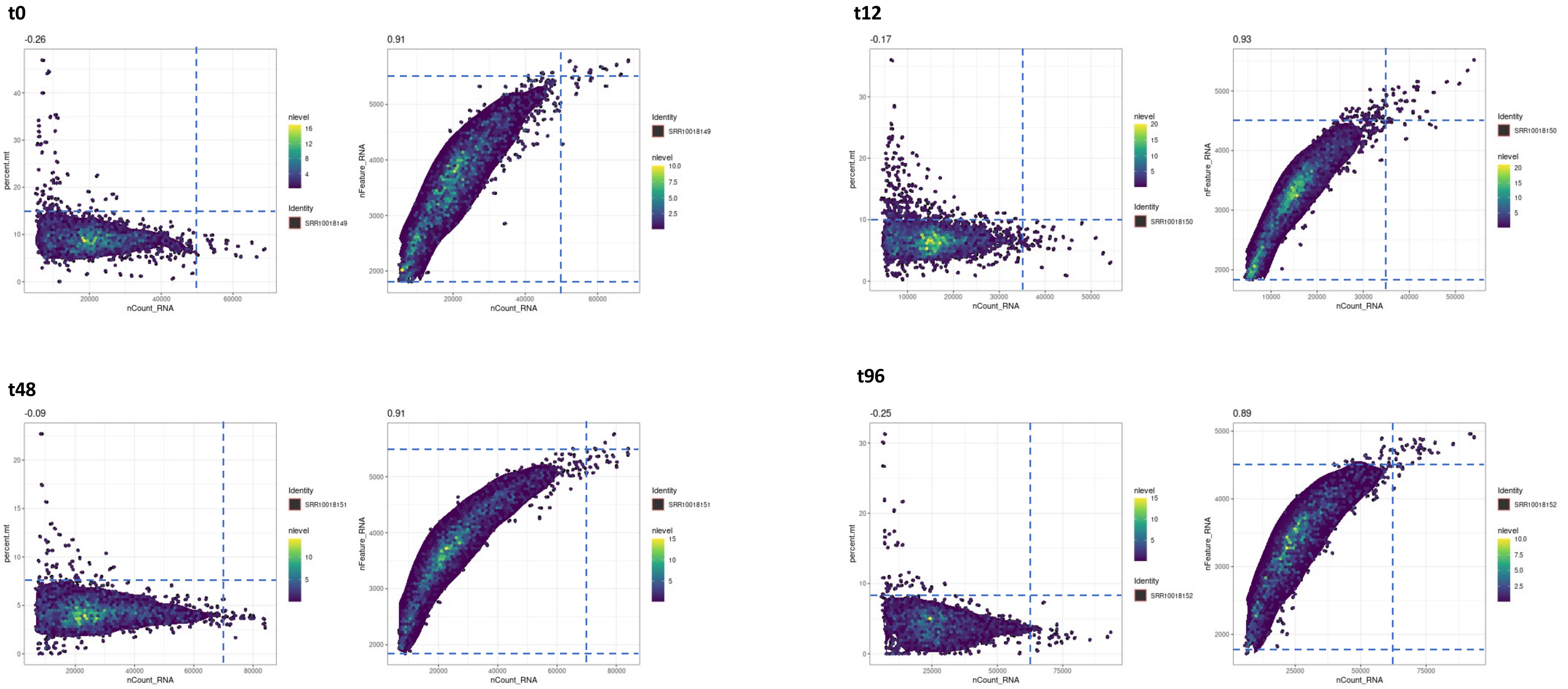


**S_Figure 4.** Density plots showing the distribution of cells based on proportion of transcripts of mitochondrial origin (left), and number of genes (right) plotted against the counts of sequencing reads in the 4 time-ponts after MCF7 drug treatment. The dotted line indicates the selected QC thresholds: mitochondrial gene expression above between 7.5 and 15%, and number of genes below 1000 or above between 4500 and 5500.
