## Supplementary material for "Improved SNV discovery in barcode-stratified scRNA-seq alignments": S_Figure 5

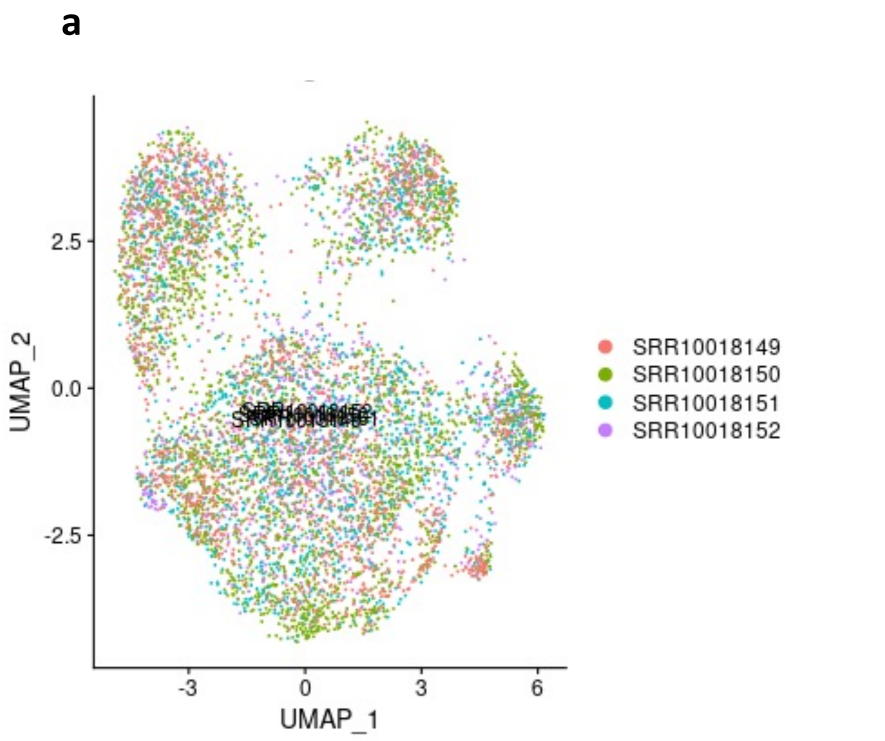


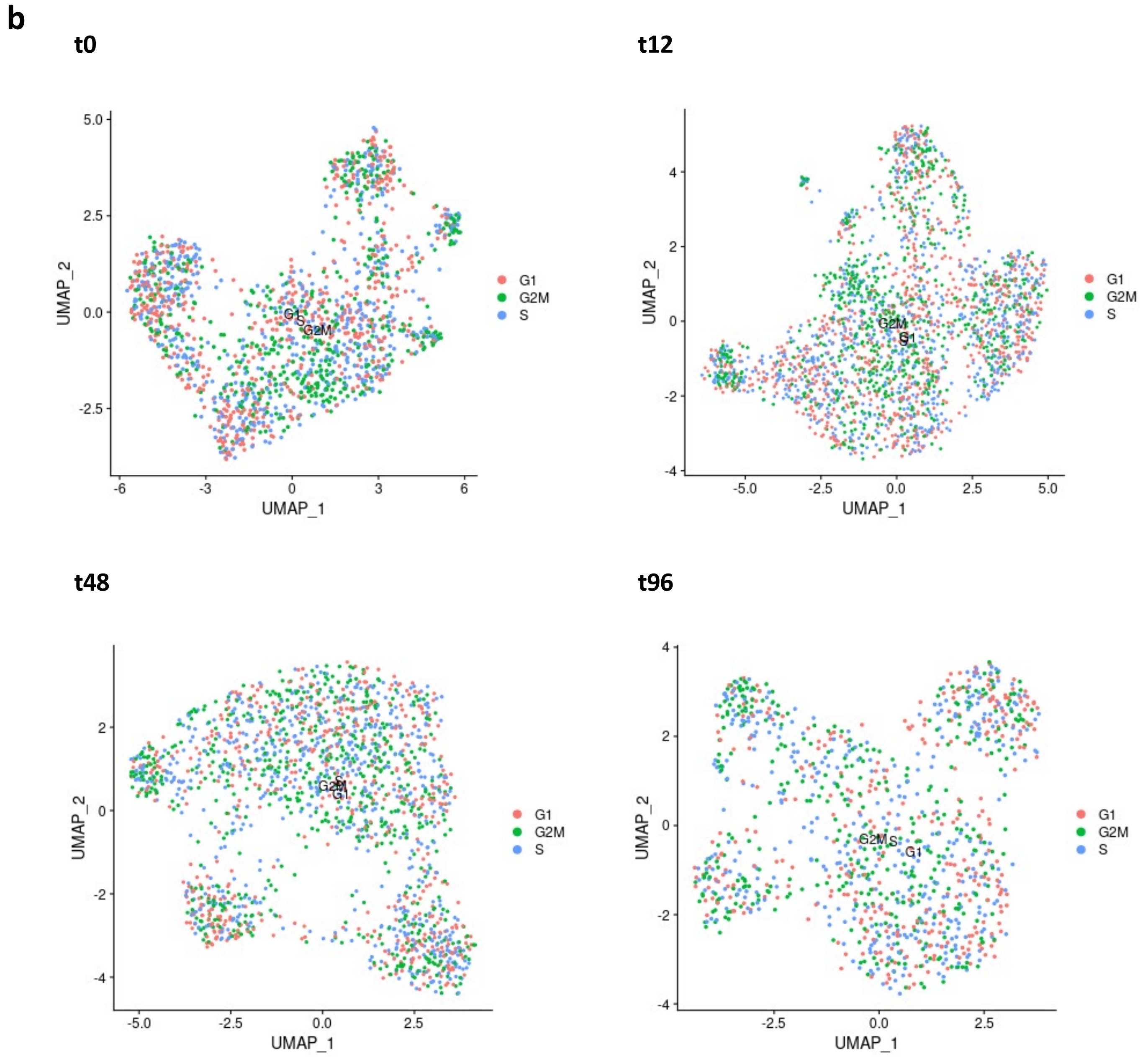


**S_Figure 5.** Two-dimensional UMAP projections of the scRNA-seq data from the 4 time-points after MCF7 drug treatment for assessment of batch and cell-cycle effects. a) No batch effects are seen in the combined t0-t96 UMAP projection. b) Cell cycle scores computed for the 4 time-points after MCF7 drug based on expression of G2/M and S phase markers assigned using Seurat after regressing out the cell cycle source of heterogeneity.
