## Supplementary material for "Improved SNV discovery in barcode-stratified scRNA-seq alignments": S_Figure 6

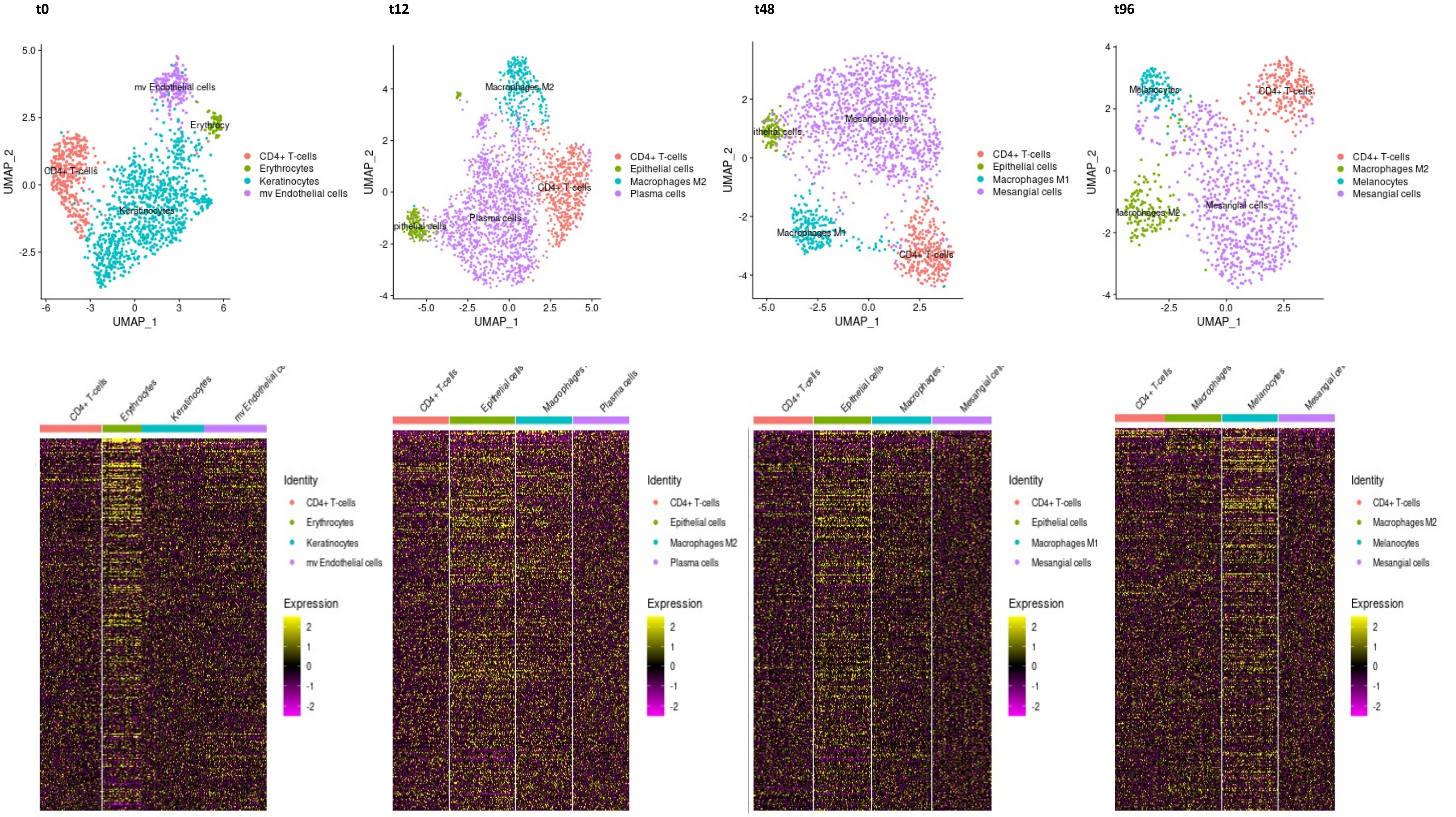


**S_Figure 6.** SingleR analysis of the data from the 4 time-points after MCF7 drug treatment. Top: SingleR Cluster Labels showing highest similarity with. Across the 4 datasets, cells with gene expression to the following cell types was found: CD4+ T-cells, Epithelial Cells, Macrophages, Endothelial cells, Erythrocytes, Keratinocytes, Plasma cells, and Mesanglial cells. Bottom: SingleR gene expression heat maps.
